# N-terminal autoprocessing and acetylation of MARTX Makes-Caterpillars Floppy-like effector is stimulated by ADP-Ribosylation Factor 1 in advance of Golgi fragmentation

**DOI:** 10.1101/736421

**Authors:** Alfa Herrera, John Muroski, Ranjan Sengupta, Hong Hanh Nguyen, Shivangi Agarwal, Rachel R. Ogorzalek Loo, Seema Mattoo, Joseph A. Loo, Karla J. F. Satchell

**Affiliations:** Department of Microbiology-Immunology, Northwestern University Feinberg School of Medicine, Chicago, IL 60611, USA; Department of Chemistry and Biochemistry, University of California-Los Angeles, Los Angeles, CA, 90095, USA; Department of Biological Sciences, Purdue University, West Lafayette, IN, 47907, USA; Purdue Institute for Inflammation, Immunology and Infectious Diseases, Purdue University, West Lafayette, IN, 47907, USA; Department of Biological Chemistry, David Geffen School of Medicine, UCLA Molecular Biology Institute, and UCLA/DOE Institute of Genomics and Proteomics, University of California-Los Angeles, Los Angeles, CA, 90095, USA

## Abstract

Studies have successfully elucidated the mechanism of action of several effector domains that comprise the MARTX toxins of *Vibrio vulnificus*. However, the biochemical linkage between the cysteine proteolytic activity of Makes Caterpillars Floppy-like (MCF) effector and its cellular effects remains unknown. In this study, we identify the host cell factors that activate *in vivo* and *in vitro* MCF autoprocessing as ADP-Ribosylation Factor 1 (ARF1) and ADP-ribosylation Factor 3 (ARF3). Autoprocessing activity is enhanced when ARF1 is in its active (GTP-bound) form compared to the inactive (GDP-bound) form. Subsequent to auto-cleavage, MCF is acetylated on its exposed N-terminal glycine residue. Acetylation apparently does not dictate subcellular localization, as MCF is found localized throughout the cell. However, the cleaved form of MCF gains the ability to bind to the specialized lipid phosphatidylinositol 5-phosphate enriched in Golgi and other membranes necessary for endocytic trafficking, suggesting a fraction of MCF may be subcellular localized. Traditional thin-section electron microscopy, high-resolution cryoAPEX localization, and fluorescent microscopy show that MCF causes Golgi dispersal resulting in extensive vesiculation. In addition, host mitochondria are disrupted and fragmented. Surprisingly, ARF1 is not itself processed or post-translationally modified by MCF. Further, only catalytically inactive MCF stably associates with ARF1, thus serving as a substrate trap. Our data indicate that ARF1 is a cross-kingdom activator of MCF, but reveal that MCF mediates cytotoxicity likely by directly targeting another yet to be identified protein. This study begins to elucidate the biochemical activity of this important domain and gives insight into how it may promote disease progression.

## 1. INTRODUCTION

*Vibrio vulnificus* is a Gram-negative pathogen found in warm marine environments that is capable of causing life-threatening gastrointestinal and wound infections, particularly from handling or consumption of seafood. These infections are rare, although the incidence of *V. vulnificus* is increasing in the US, in part due to expanding geographical distribution due to warming seawaters (Baker-Austin et al., 2012; Deeb, Tufford, Scott, Moore, & Dow, 2018; King et al., 2019). The rising incidence of *V. vulnificus* infections is extremely concerning given its severity, as it often leads to fatal sepsis (Horseman & Surani, 2011). About 90% of patients are hospitalized and more than 50% die, often as fast as 48 hours after symptom onset (CDC, 2014; Gulig, Bourdage, & Starks, 2005). These infections are costly due to high mortality rates and negative impact on economies that depend on recreation and seafood harvesting (Hoffman & Batz, 2015; Morgan, 2010). Currently in the US, *V. vulnificus* accounts for only 0.001% of infections, but 3% of all food-related deaths (Mead et al., 1999). According to the US Department of Agriculture, *V. vulnificus* exerts the highest per-case economic burden of any food-borne disease and its total burden exceeds *Shigella* and *Escherichia coli* O157:H7 (Hoffman & Batz, 2015).

For both wound and intestinal infections, the primary *V. vulnificus* virulence factor associated with sepsis and subsequent death is the composite multifunctional-autoprocessing repeats-in-toxins (MARTX) toxin encoded by the gene *rtxA1* (Satchell, 2015). Expression of the MARTX toxin can increase lethality of *V. vulnificus* up to 2600-fold (Kwak, Jeong, & Satchell, 2011). The *V. vulnificus* MARTX toxin is secreted from the bacterium as a single large polypeptide. It is composed of conserved glycine-rich repeats at both the N- and C-termini between which there are variable effector domains and a cysteine protease domain (CPD). The repeat regions are thought to recognize eukaryotic host cells via a yet undetermined mechanism and then form a pore in the plasma membrane. The effectors and CPD are then translocated across the membrane through this pore. Within the cell, the CPD binds inositol hexakisphosphate, which activates the CPD to auto-cleave at multiple sites to release each of the arrayed effectors individually into the host cytoplasm. The MARTX toxin thus serves as an effector delivery platform for translocation of multiple toxic virulence factors inside cells as a single bolus (Gavin & Satchell, 2015; Kim, 2018; Satchell, 2015).

Across distinct *V. vulnificus* isolates, the effector content can vary from 2 to 5 effectors selected from a total of nine known effector domains (Kwak et al., 2011; Ziolo et al., 2014). The effector repertoire can be exchanged by horizontal gene transfer and recombination events such that even closely related strains can have different MARTX toxin variants (Roig, Gonzalez-Candelas, & Amaro, 2011). The effector domain most commonly present in *V. vulnificus* MARTX toxins is the Makes Caterpillars Floppy-like (MCF) effector domain (Agarwal, Agarwal, Biancucci, & Satchell, 2015; Agarwal, Zhu, Gius, & Satchell, 2015). It is found in all Biotype 1 and 2 strains that cause human infections and is even duplicated in the MARTX toxin of some strains (Kwak et al., 2011; Roig et al., 2011).

The MCF effector domain is 376 aa in size. In representative strain CMCP6, MCF is encoded by gene *rtxA1* (*vv2_0479*) nucleotides 9610-10737 to comprise residues 3204-3579 of the full-length MARTX toxin. Once within cells, MCF has been shown to induce the intrinsic apoptotsis pathway, resulting in cell rounding, vesiculation, and loss of cell proliferation (Agarwal, Agarwal, et al., 2015; Agarwal, Zhu, et al., 2015). MCF has been shown to activate caspase-9, 7, 3 and PARP-γ, and to induce nuclear shrinkage and fragmentation. It also upregulates the proapoptotic proteins Bax, Bak, and Bad to induce loss of mitochondrial transmembrane potential resulting in increased release of cytochrome c into the cytosol.

Sequence homology places MCF into the C58 family of cysteine peptidases (Rawlings, Waller, Barrett, & Bateman, 2014). Site-directed mutagenesis revealed that the conserved Cys-148 residue (aa 3351 in full-length toxin) is essential for MCF cytotoxicity. However, even though other putative catalytic residues His-55 (aa 3258 in full-length toxin) or Asp-142 (aa 3345 in full-length toxin) are conserved by sequence alignment with the C58 peptidases that use a Cys-His-Asp (CHD) catalytic triad, alanine substitution of these residues reduced, but did not eliminate, cell rounding (Agarwal, Agarwal, et al., 2015). Instead, MCF has an Arg-Cys-Asp (RCD) peptide triad at residues 147-149 (aa 3350-3352 in full-length toxin) that is necessary for its cytotoxic effects. A homology search revealed numerous known and putative toxin proteins that contain domains with sequence similarity to MCF that also contain the RCD/Y motif (Agarwal, Agarwal, et al., 2015). We suggest these proteins form a new subfamily of the C58 peptidases. Thus far, no structures of a peptidase from this family has been determined so the structural implications of the RCD/Y motif compared to the CHD motif remain unknown.

Upon ectopic expression in host cells, MCF undergoes autoprocessing at its N-terminus cleaving itself between Lys-15 and Gly-16 dependent upon an intact RCD motif (Agarwal, Agarwal, et al., 2015). This autoprocessing could be reproduced with recombinant MCF, but only upon addition of eukaryotic cell lysate to the reaction. In this study, we identify the cross-kingdom autoprocessing-stimulating host factor as Class I and II ADP-ribosylation factors (ARFs), and most prominently ARF1. We further show that subsequent to autoprocessing, the exposed N-terminal Gly-16 is acetylated. MCF then also gains the ability to bind phosphatidylinositol-5-phosphate (PtdInsP5), a lipid enriched on the Golgi apparatus. The expression of MCF in cells then leads to increased vesiculation in the cells caused by dissolution of the Golgi apparatus. Destruction of the Golgi, and also the mitochondria, suggests a mechanism for induction of caspases leading to MARTX toxin dependent cellular apoptosis (Lee, Choi, & Kim, 2008).

## 2. RESULTS

### 2.1. Purified ARF1 and ARF3 GTPases induce autoprocessing of recombinant MCF protein

MCF is known to be induced to auto-cleave by an unidentified eukaryotic host cell factor. To identify this host factor, MCF encoding gene sequences were truncated to mimic the cleaved form of MCF (cMCF, aa 16-376) and then modified to alter the codon for the catalytic Cys-148 to alanine (cMCF^CA^). The modified gene was ectopically expressed in human embryonic kidney (HEK) 293T cells fused with an N-terminal FLAG and C-terminal hemagglutinin (HA) tag to express dual-tagged cMCF^CA^ (dt-cMCF^CA^). Anti-HA immunoprecipitation revealed a protein of approximately 20 kDa that bound to dt-cMCF^CA^ (Fig 1A). Peptide sequencing of the excised band revealed nine unique peptides. Seven mapped to the human protein ADP-Ribosylation Factor (ARF1) and the nearly identical protein ADP-Ribosylation Factor 3 (ARF3). One peptide matched exclusively to ADP-Ribosylation Factor (ARF4) and another to ADP-Ribosylation Factor 6 (ARF6) (Fig 1B).

**Figure 1.**
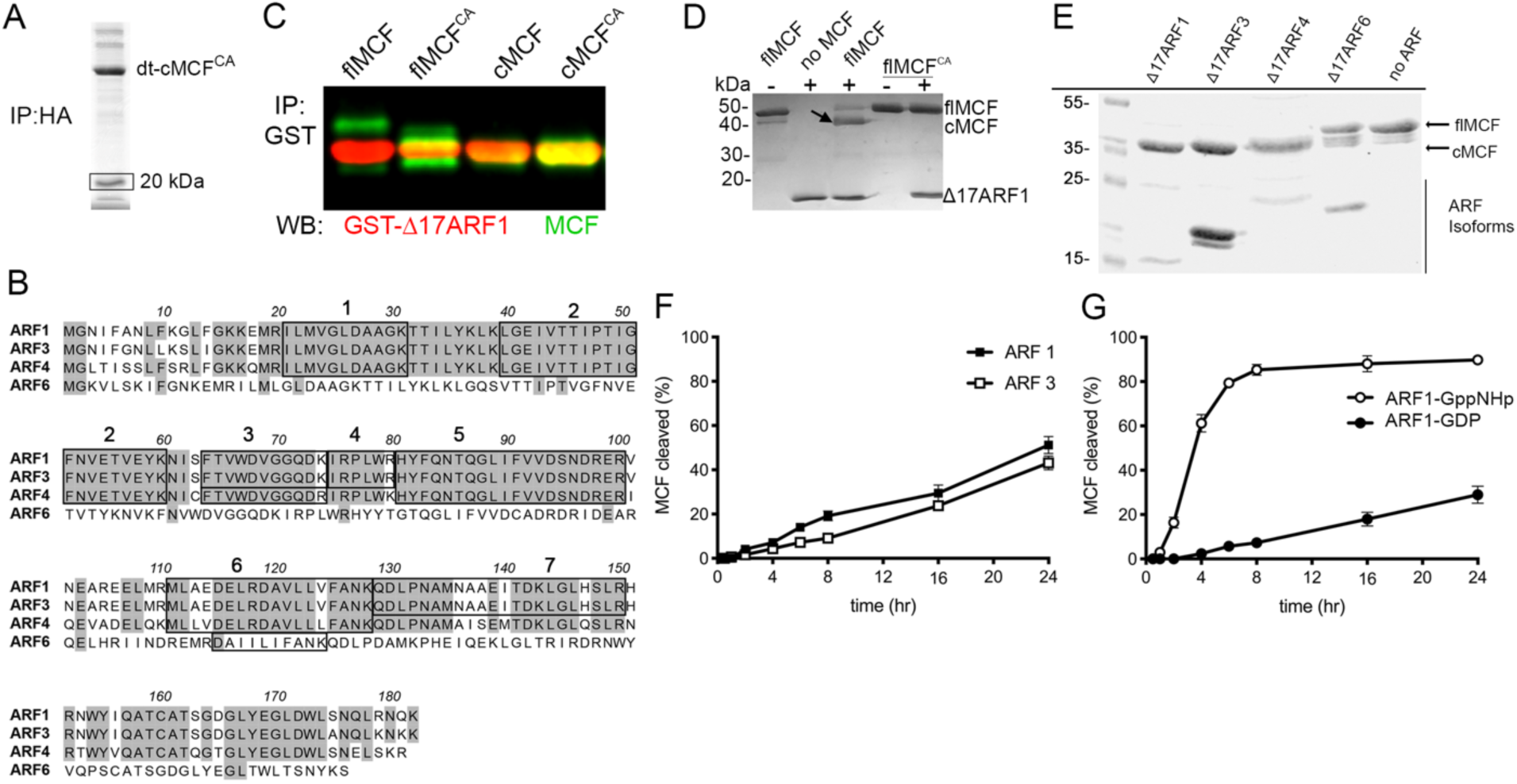
MCF autoproteolytic cleavage is induced by ARF GTPases. A. dt-cMCF^CA^ was immunoprecipitated (IP) from HEK293T cell lysate using anti-HA antibody. Box indicates band excised for mass spectrometry peptide sequencing. B. Amino acid sequence alignment of major isoforms of ARFs with boxes indicating identified peptides. C. Purified GST-Δ17 ARF1 was incubated with either MCF, MCF^CA^, cMCF, or cMCF^CA^ at 37°C. Western blot on samples following anti-GST IP using anti-MCF (green bands) and anti-ARF1 (red bands). showing catalytically active and inactive flMCF and cMCF (green bands) can bind ARF1(red bands) *in vitro*. D. Recombinant flMCF and flMCF^CA^ incubated with ΔARF1 indicate induced autoprocessing is dependent on MCF catalytic active site. E. Auto-cleavage induced by purified ARF isoforms incubated with MCF for 24 hours. F, G. Autoprocessing of flMCF at 37°C for time indicated induced by full-length ARF1 or ARF3 (F) or with ARF1 pre-loaded with GDP or non-hydrolyzable GTP (GppNHp) (G). Representative gels (*n*>3), with percent cleaved MCF was determined by densitometry of bands on Coomassie stained gels (%=(cMCF/(cMCF+flMCF)*100 from three independent reactions.

To confirm identification of ARF1 as a host factor that induces autoprocessing, full-length MCF (flMCF) with and without its catalytic Cys-148 (flMCF^CA^), cleaved MCF (cMCF), and catalytically inactive cleaved MCF (cMCF^CA^) were purified as 6xHis-tagged recombinant proteins. Truncated Δ17ARF1 fused with glutathione-S-transferase (GST-Δ17ARF1) was purified as removal of the flexible N-terminus is known to increase protein solubility (Kahn et al., 1992). Both catalytically active and inactive flMCF and cMCF forms co-precipitated with GST-Δ17ARF1 confirming that MCF and ARF1 interact and that the binding does not involve the flexible N-terminus of ARF1 (Fig 1C).

The flMCF proteins were next incubated with Δ17ARF1 purified as a 6x-tagged recombinant protein to test stimulation of autoprocessing. Δ17ARF1 stimulated flMCF cleavage and the processing was autocatalytic as flMCF^CA^ was not processed even in the presence of Δ17ARF1 (Fig 1D). Δ17ARF3 and Δ17ARF4 also stimulated autoprocessing, while Δ17ARF6 did not (Fig. 1E). Full-length ARF1 and ARF3 with the N-terminus intact were also able to stimulate autoprocessing with similar kinetics (Fig. 1F).

ARFs are all members of the Ras superfamily of small GTPases that cycle between an active (GTP-bound) and inactive (GDP-bound) form. ARF1 pre-loaded with GTP-analog GppNHp induced more rapid and more complete MCF auto-cleavage (85 +/-3%) compared to ARF1 pre-loaded with GDP (29 +/- 13%) (Fig 1G).

Altogether, these data demonstrate that ARF1 is a host factor that binds and stimulates autoprocessing of MCF. The nearly identical protein Class I ARF3 and the Class II ARF4 can also serve as cross-kingdom host factors capable of inducing MCF auto-cleavage. Further, there is a strong preference for induction of autoprocessing by the active (GTP-bound) form of ARFs.

### 2.2. MCF^CA^ functions as a trap for ARF1 in cells

To identify which of the major ARF isoforms are important within cells for induced autoprocessing, dt-cMCF^CA^ was co-expressed in HEK293T cells with Myc-tagged ARF proteins. ARF1, but not ARF4 or ARF6, co-immunoprecipitated with cMCF^CA^ (Fig 2A). We were unable to reproducibly pulldown ARF3 with dt-cMCF^CA^ across independent experiments and thus were not able to confirm this interaction. Thus, although multiple ARF isoforms stimulate MCF autoprocessing *in vitro*, ARF1 is the only isoform that consistently associates with dt-cMCF^CA^ in cells.

**Figure 2.**
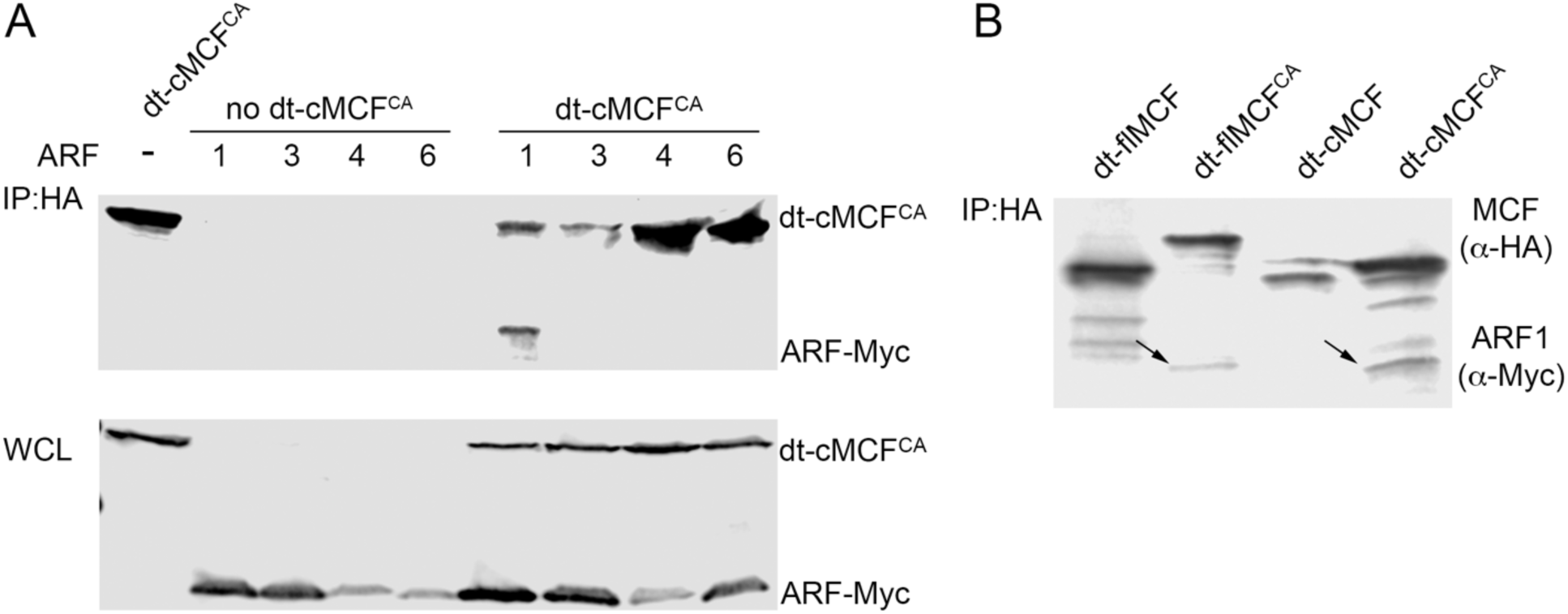
MCF^CA^ stably associates ARF1 in HEK293T cells. A. HEK293T cells were co-transfected with dt-cMCF^C148A^ and either Myc-tagged ARF1, 3, 4, or 6. Western blot of 40 µg of total protein in whole cell lysates (bottom) recovered from cells. Western blot of 600 µg of total protein from whole cell lysate after IP using anti-HA beads (top). MCF and ARFs detected by anti-HA and anti-Myc antibodies respectively with representative gel shown (n>3). B. HEK293T cell were co-transfected with ARF1-Myc and either dt-flMCF, dt-flMCF^CA^, dt-cMCF, or dt-cMCF^CA^. Western blot on cell lysates recovered from these cells following anti-HA IP as in (A).

Consistent with its function as a host factor for induction of autoprocessing of flMCF, ARF1 was also co-immunoprecipitated with dt-flMCF^CA^, indicating interaction between the two proteins occurs prior to the autoprocessing event. By contrast, ARF1 did not co-immunoprecipitate with either dt-flMCF or dt-cMCF if the catalytic Cys-148 was not modified to alanine (Fig 2B). This is not due to an inability of the proteins to associate because, as shown above, GST-Δ17ARF1 co-precipitated with both catalytically active and inactive flMCF and cMCF *in vitro* (Fig 1C). These data indicate that, in cells, MCF autoprocessing results in release and/or loss of the ability to bind to ARFs such that only catalytically inactive MCF functions as a protein trap. Further, since overexpression or delivery of catalytically inactive MCF to cells does not itself cause cytotoxicity (Agarwal, Agarwal, et al., 2015; Agarwal, Zhu, et al., 2015), trapping of ARFs is not sufficient for MCF-mediated cytotoxicity.

Repeated attempts to identify an MCF-specific processing or post-translational modification of ARF1 or ARF3 *in vitro* or on proteins recovered from cells were not successful. We did not observe any visible changes to the size of ARF1-Myc when co-transfected with either full length or cMCF or the catalytically inactive versions by western blot (Fig 2A-B). We also did not identify any MCF-specific post-translational modifications on ARF1 or ARF3 using a bottom-up proteomics strategy (Supplemental Fig 1). Therefore, it is possible that while ARF1 can induce MCF to autoprocess, MCF-mediated processing or covalent modification of ARFs is not the mechanism of cell cytotoxicity. Thus, we speculate the preferred association of MCF with ARF1-GTP functions to localize MCF to a proper membrane where it is activated by autoprocessing, but then released from ARF1, possibly to access other targets.

### 2.3. After autoprocessing, MCF is N-terminally acetylated

In cells, MCF undergoes additional modifications. Previously, attempts to identify the autoprocessing site on MCF from protein recovered from cells revealed the N-terminus was blocked to Edman degradation. The N-terminal block does not occur *in vitro*. The Edman degradation analysis of immunoprecipitated MCF was repeated for this study and confirmed to be blocked (Supplemental Fig 2A and 2B). Further, two-dimensional gel analysis of MCF expressed in cells showed that dt-flMCF undergoes *in vivo* autoprocessing to release its N-terminal FLAG tag, coincident with a shift in electrophoretic mobility, which was detected by western blotting using anti-HA antibody directed against the C-terminal tag. It was further noted that processed flMCF separated into discrete populations that differ in isoelectric point. Catalytically inactive dt-flMCF^CA^ does not shift in size, consistent with the autoprocessing being autocatalytic, or separate into different populations (Fig 3A).

**Figure 3.**
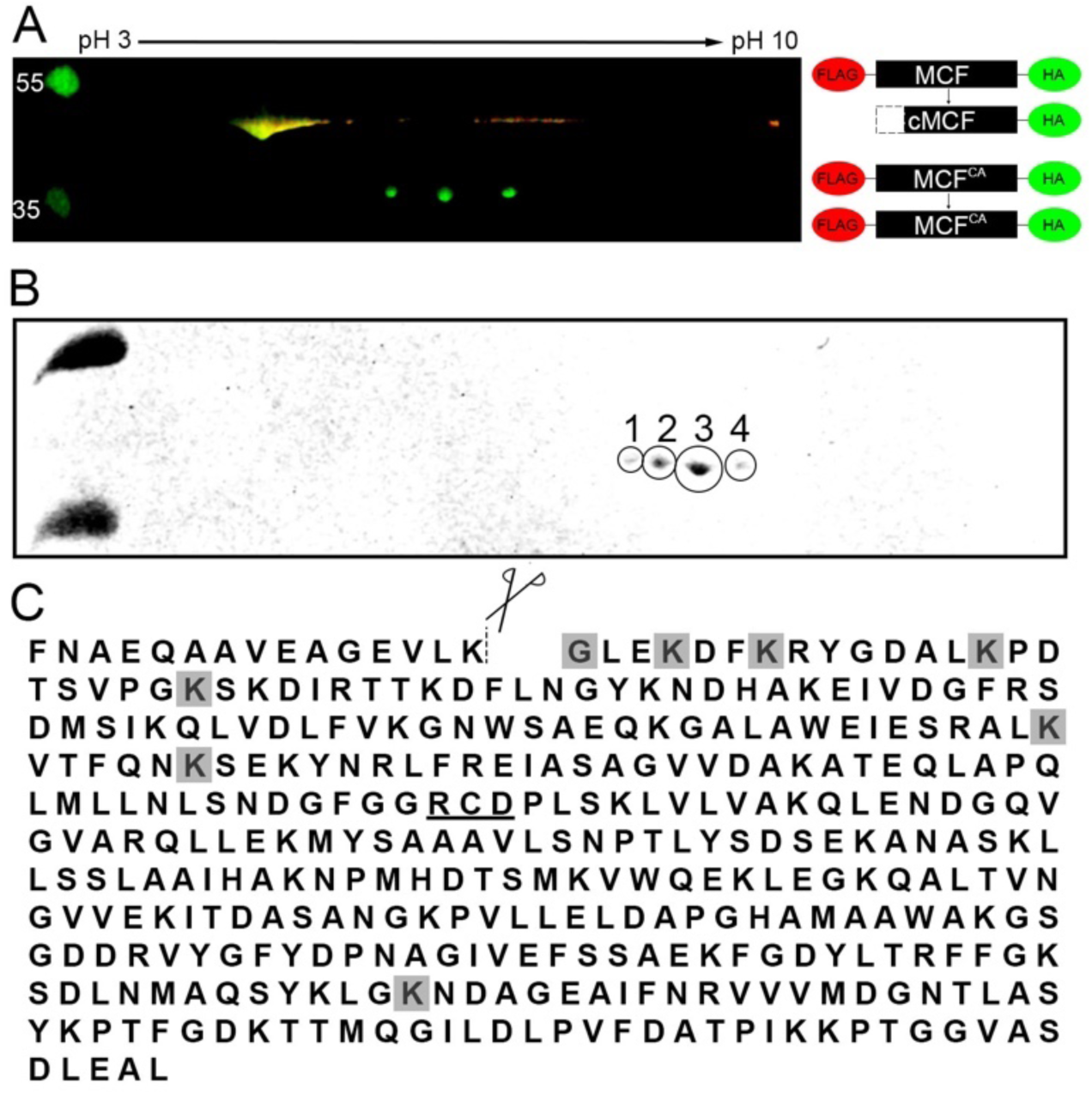
MCF is N-terminally acetylated following auto-cleavage. A. Anti-HA IP on cell lysates of HEK293T cells ectopically expressing dt-MCF or dt-MCF^CA^ was individually completed. Samples were then combined for two-dimensional western analysis using IPG strips pH 3-10 with western blot completed using anti-FLAG (red) and anti-HA (green) antibodies. B. Two-dimensional analysis with Coomassie gel was completed using dt-MCF as in (A). Gel shows populations that were individually excised and analyzed by bottom up mass spectrometry. C. Protein sequence of MCF designating auto-cleavage site (dashed-line), and catalytic residues important for auto-proteolytic activity (underlined). Shaded residues are those found to be acetylated in populations 1-4.

To identify potential post-translational modifications on MCF, the four individual populations were excised and analyzed by bottom-up mass proteomics (Fig 3B); the analysis revealed that Gly-16, which is exposed following auto-cleavage, was acetylated (Fig 3C and Supplemental Fig 3). The acetylation was identified in populations 2 and 3, but not population 1 or 4 (Supplemental Fig 3). Acetylation of other lysine residues in populations 2, 3, and 4 was also observed, but the modified lysine varied across different experiments or replicates of the same population (Fig 3C and Supplemental Fig 3). These observations indicate that only the acetylation of the N-terminal glycine of MCF is specific, while other acetylation events are stochastic modifications that likely arise from overexpression of a protein with many surface exposed lysine residues.

### 2.4. MCF binds Golgi enriched lipids and causes Golgi dispersion

ARF1 and ARF3 most commonly localize to the Golgi apparatus (Hofmann & Munro, 2006; Tsai, Adamik, Haun, Moss, & Vaughan, 1993) and ARF association with the Golgi is dependent upon its activation by GTP. As MCF preferentially associates with active ARFs, it may facilitate targeting to subcellular organelles. In addition, many proteins are directed to subcellular organelles by binding specific lipids. Lipid overlay assays show cMCF, but not flMCF, preferentially binds PtdIns5P (Fig 4). PtdIns5P is not well described but has a suggested role in membrane trafficking and is enriched in Golgi membranes (Backer, 2010). Thus GTP-bound ARFs may bring MCF to the Golgi for activation, but cMCF might be retained at the Golgi even after dissociation from ARFs due to a gain of function for PtdIns5P binding.

**Figure 4.**
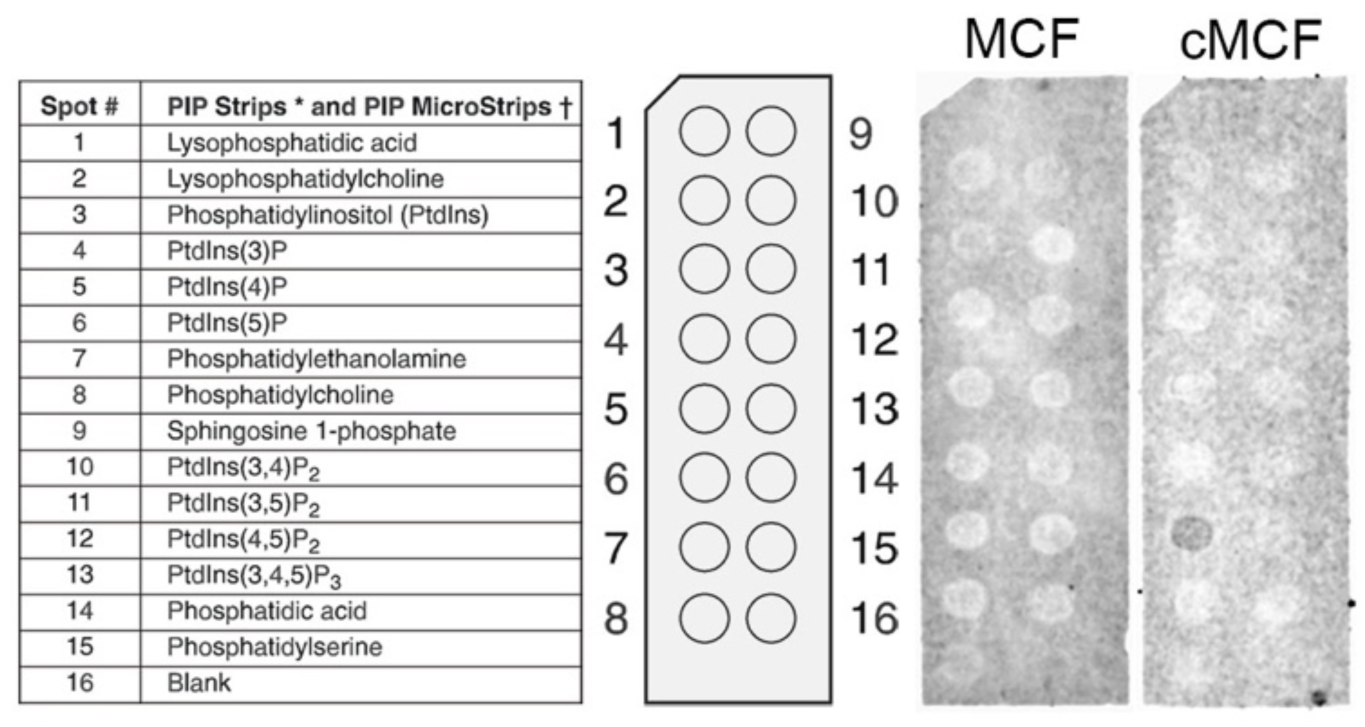
MCF associates with Golgi associated lipids. Lipid overlay assay was conducted using PIP Strips^™^ with either purified flMCF or cMCF. Blot shows cMCF binds phosphatidylinositol-5-phosphate.

The co-localization of MCF with the Golgi was thus assessed by immunofluorescence (IF). In cells transfected to express only green fluorescent protein (eGFP), the human Golgi matrix protein GM130, a marker for the *cis*-Golgi, showed the Golgi apparatus as tight and condensed at its expected position near the nuclei (Fig 5A). Catalytically inactive MCF^CA^ frequently co-localized with GM130, although protein also diffused throughout the cell (Fig 5A).

**Figure 5.**
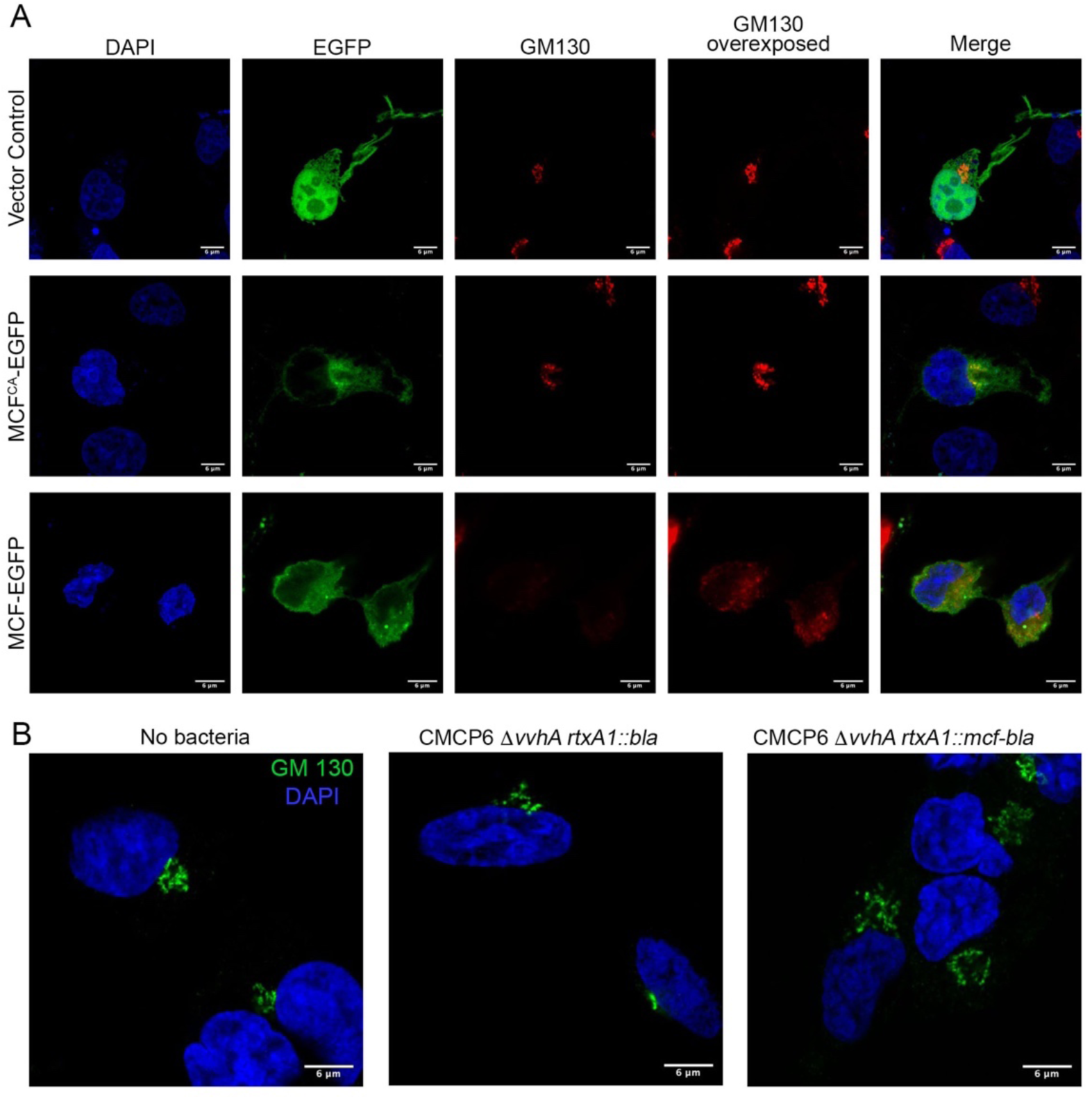
MCF induces Golgi dispersion. A. Cos7 cells were transfected with empty vector, MCF^CA^-EGFP, or MCF-EGFP (green) for 18 hours, fixed, and stained for DAPI (blue), and *cis*-Golgi marker (GM130) (red). B. Cos7 cells were infected with CMCP6 *rtxA1::bla* or CMCP6 *rtxA1::mcf-bla* at a MOI of 5 for an hour, fixed, and stained for DAPI (blue), and *cis*-Golgi marker (GM130) (green).

However, the co-localization of MCF^CA^ does not inform about possible localization of MCF after dissociation from ARFs. Thus, we examined if catalytically active MCF retains association with the Golgi or other membranes. While the Golgi in cells expressing flMCF-eGFP was disrupted and dispersed, attempts to co-localize MCF were not conclusive. Indeed, GM130 in these cells was not visible unless the image was overexposed (Fig 5A). Similar dispersion was not observed for the endoplasmic reticulum (ER)-associated protein calreticulin in cells transfected to express MCF-eGFP (Supplemental Fig 4). Thus, catalytically active MCF was found to disrupt the Golgi of host cells and the resulting damage prevented resolution of its intracellular localization using IF.

### 2.5. Golgi dispersion and mitochondrial fragmentation occurs when MCF is delivered to cells by the bacterium

To ensure Golgi disruption is not an artifact of overexpression of MCF by transient transfection, cells treated with live *V. vulnificus* were imaged. Sequences corresponding to the MARTX toxin effectors had been previously deleted from the *rtxA1* gene in strain CMCP6 strain and replaced with in-frame beta-lactamase gene sequences to generate a fusion gene *rtxA1::bla* that encodes an effector-less MARTX toxin but delivers beta-lactamase as a marker for intact protein and translocation function. Sequences for MCF were then introduced to generate *rtxA1::mcf-bla* that expresses a MARTX toxin that delivers only the MCF effector (Agarwal, Zhu, et al., 2015).

Treatment of cells with CMCP6 *rtxA1::bla* showed the Golgi remained perinuclear and condensed similar to non-intoxicated cells. In contrast, cells intoxicated with CMCP6 *rtxA1::mcf-bla* showed Golgi was dispersed and spreading away from the nucleus (Fig 5B). In addition, the impact of MCF on mitochondria was also visualized using MitoTracker Green FM. The mitochondrial network of cells treated with the control strain CMCP6 *rtxA1::bla* appeared as long dynamic fragments throughout the cell (Fig 6). In contrast, the mitochondria of cells intoxicated with CMCP6 *rtxA1::mcf-bla* were shortened and fragmented in comparison. The mitochondrial network in these cells was composed of more short round fragments than long branches (Fig 6).

**Figure 6.**
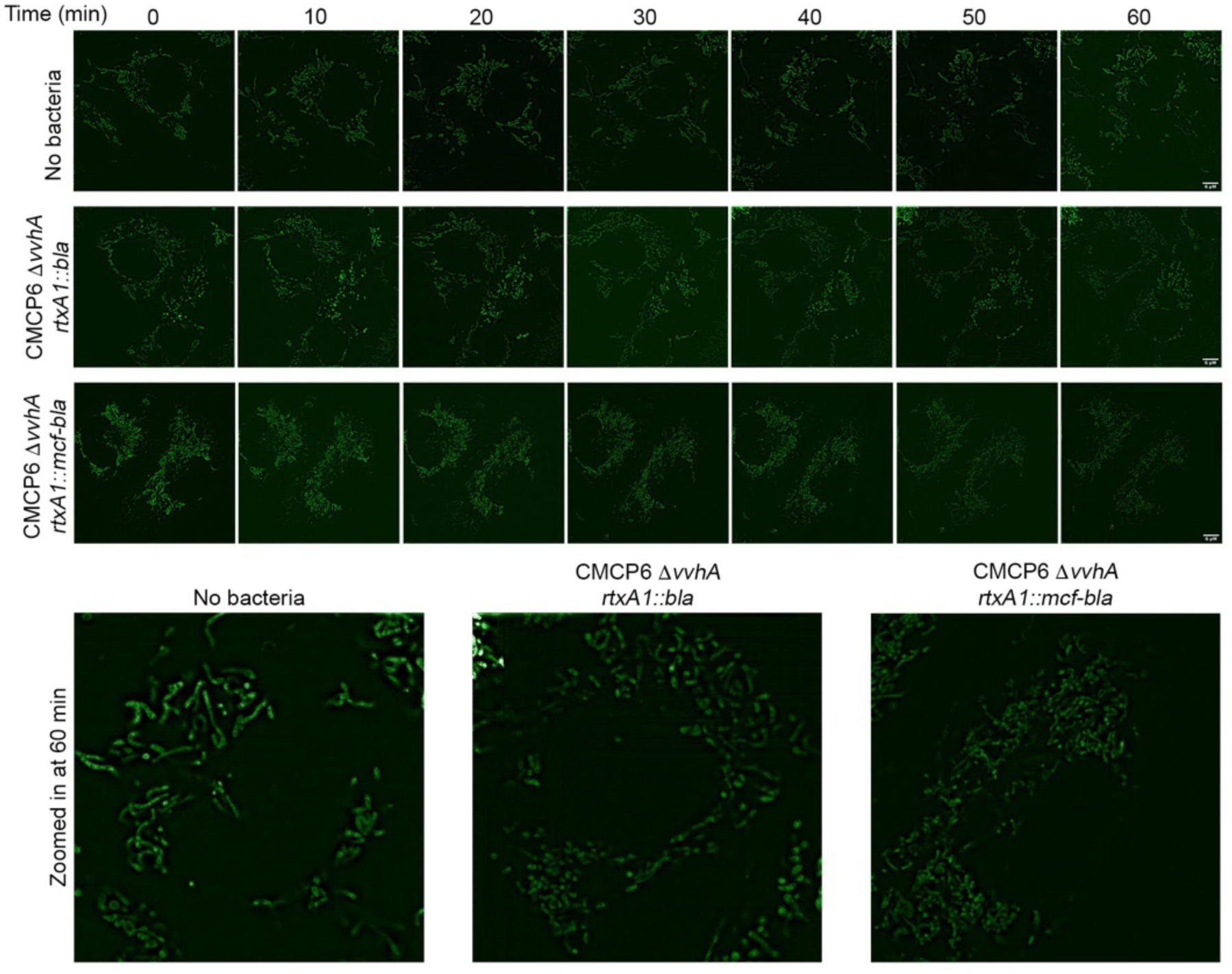
MCF disrupts the mitochondrial network. A. Cos7 cells were incubated with MitoTracker™ Green FM to visualize mitochondria prior to infection with CMCP6 *rtxA1::bla* or CMCP6 *rtxA1::mcf-bla* at a MOI of 5 for an hour. Images taken prior to intoxication and every 10 minutes after. Insets show enlarged image of one cell for each condition at 60 minutes.

These data show that MCF delivered to cells by bacteria induces intracellular damage of both the Golgi and the mitochondria.

### 2.6. Transmission electron microscopy shows MCF induces extensive vesiculation of the Golgi apparatus

To gain an ultrastructural understanding of how catalytically active MCF induces Golgi dispersion, transmission electron microscopy was employed. As expected, at 10 hours after transfection, PBS mock transfected HeLa cells showed a Golgi with typical stacked morphology (Fig 7A, green G). By contrast, in cells ectopically expressing MCF, there are a significant number of vesicles (Fig 7B, purple V) in the cytoplasm localized near the Golgi, with some enmeshed within the Golgi complex (Fig 7B, Supplemental Fig 5A). At higher magnification, it was evident that, although some of the Golgi retained a classical stacked morphology some of the cisternae appeared herniated transitioning into a more vesicular morphology (Fig 7B, Supplemental Fig 5A). Additionally, an abundance of autolysosomes was also observed. Some autolysosomes contained vesicles, indicating ongoing clearance off these vesicles (Fig 7E). At 15 hours post transfection, the cytoplasm was inundated with a variety of different sized pleomorphic vesicles (Fig 7D, Supplemental Fig 5B). In these cells the typical stacked structure of the Golgi apparatus, which is seen in mock treated cells, was not observed (Fig 7C).

**Figure 7.**
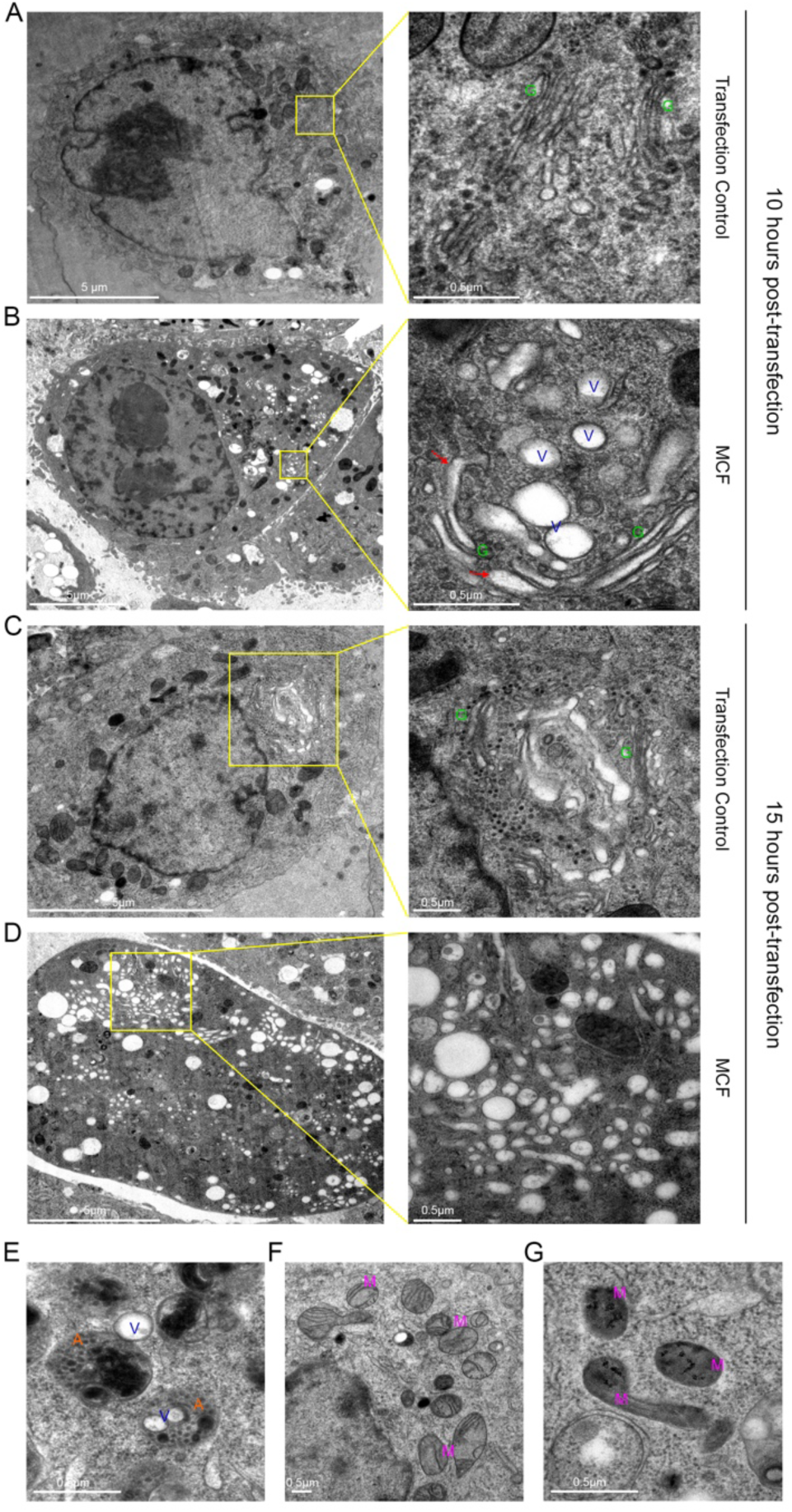
MCF causes vesiculation of the Golgi apparatus as seen by transmission electron microscopy. A-G. Thin-section electron microscopy of HeLa cells ectopically expressing MCF, or mock transfected for either 10 (A)(B) or 15 hours (C)(D) post-transfection. A-D. On right, higher magnification of boxed region on left showing Golgi (G), herniated Golgi (red arrows), and vesicles (V). E. Autolysomes (A) clearing MCF induced vesicles. F, G. Mitochondria (M) of mock transfected cells (F) and MCF transfected cells (G) at 10 hours post-transfection.

The electron microscopy data correlate with the IF observations showing darkened mitochondria at 10 hours post-transfection with condensed disintegrating cristae in cells ectopically expressing MCF (Fig 7G). This was in contrast to control cells where mitochondria have an oval shape, with visible cristae (Fig 7F). These data show that both the mitochondria and the Golgi are impacted by expression of MCF.

### 2.7. MCF localizes to the Golgi apparatus, the membrane of the vesicular structures and the cytosol

As both N-terminal acetylation and binding to PtdIns5P could dictate subcellular localization, we investigated localization of MCF using electron microscopy based high resolution cryoAPEX technology (Sengupta, Poderycki, & Mattoo, 2019). Briefly, MCF was expressed in HeLa cells as a fusion with a soybean ascorbate peroxidase (Apex2). This tag catalyzes a peroxide reaction that converts diaminobenzidine (DAB) into an insoluble osmophilic precipitate. This precipitate could then be preferentially and specifically stained with osmium tetroxide thereby revealing the subcellular localization of MCF at a nanometer-scale resolution, with MCF-Apex2 expressing cells easily distinguished from non-expressing cells due to dark staining of the cytoplasm and presence of cytoplasmic vesicles compared to the mock transfected control (Fig 8B-C, and 8A). This ubiquitous staining of the cytosol indicates a large pool of MCF is present in cytosol. Interestingly, staining was also detected on the membrane of the Golgi stacks visible in samples from the 10-hour time-point as well as on vesicles at both time points (Fig 8B-C, blue arrows). In comparison, control cells transfected with MCF-Apex2 for 10 hours not treated with H_2_O_2_ exhibit a relatively clear cytoplasm and lack membrane staining of vesicles (Fig 8D). Cells expressing MCF also had darkened staining at the nuclear membrane compared to control cells, indicating a presence of the effector here as well (Fig 8B-C).

**Figure 8.**
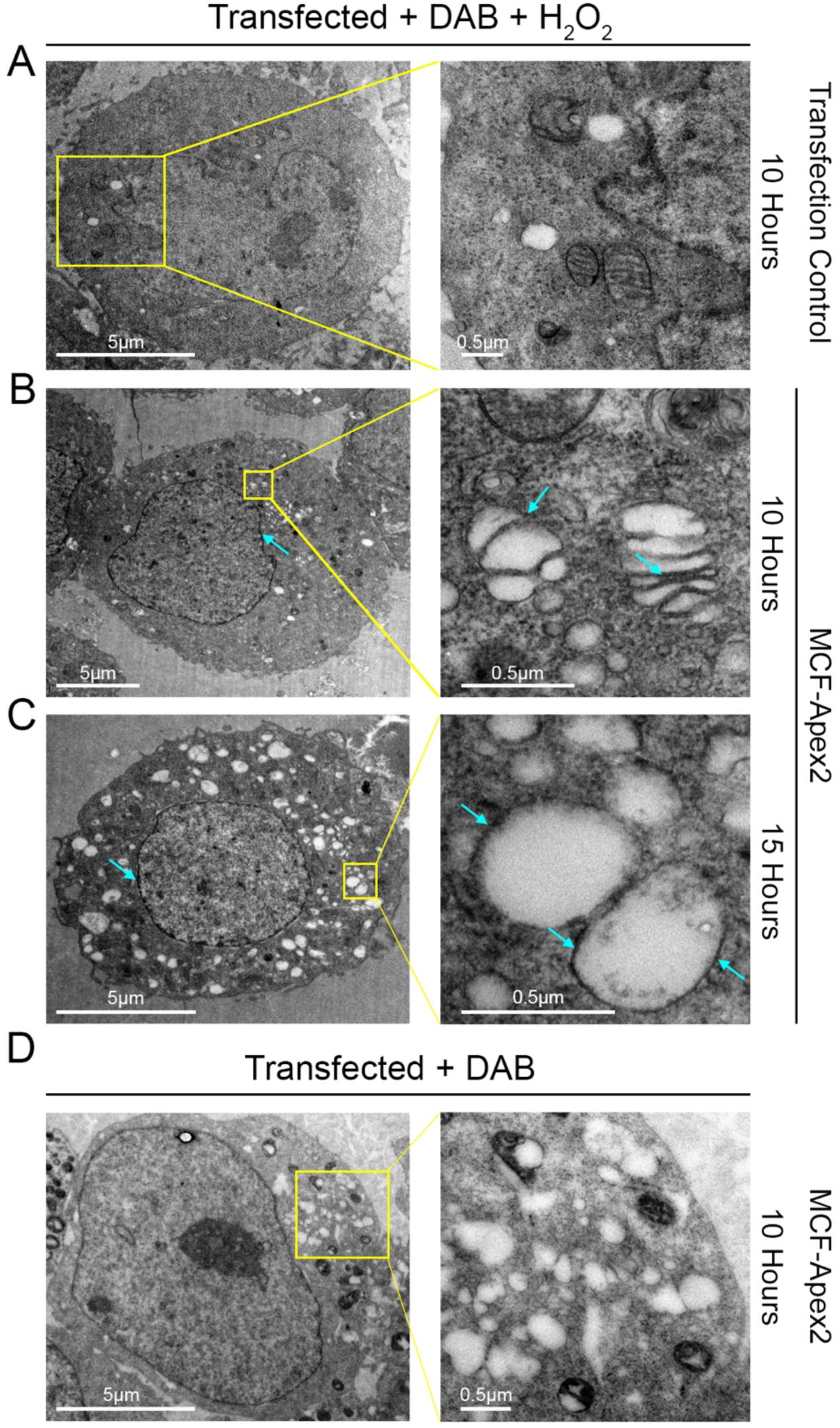
MCF concentrates in the cytoplasm, Golgi cisternae, and nuclear and vesicle membranes. A-D. Thin-section electron microscopy of resin embedded HeLa cells treated with diaminobenzidine (DAB) + H_2_O_2_ (A-C) or just DAB (D) ectopically expressing MCF (B) (C) (D), or mock transfected (A) for either 10 or 15 hours post-transfection. On right, higher magnification of boxed region on left. B-D. Light blue arrows show darkened Apex 2 staining, indicating high concentrations of MCF, seen with H_2_O_2_ treatment (B) (C) and not without (D).

## 3. Discussion

ADP-ribosylation factors (ARFs) were first identified as activators of the ADP-ribosylation activity of the bacterial protein cholera toxin (Gillingham & Munro, 2007). They subsequently have been shown to have important roles as regulators of eukaryotic cell membrane trafficking (Gillingham & Munro, 2007; Muthamilarasan, Mangu, Zandkarimi, Prasad, & Baisakh, 2016). Depending on the isoform, they localize to specific subcellular locations, including the Golgi, plasma membrane, and endosomes. The Golgi apparatus is generally composed of tight perinuclear Golgi stacks and plays a central role in membrane trafficking, wherein it receives, post-translationally modifies and sorts proteins from the ER to their appropriate destinations (Liu & Storrie, 2015; Wei & Seemann, 2010). At the Golgi, ARF1 is recognized by its guanine nucleotide exchange factors (GEF), inducing a conformational change that exposes its N-terminal amphipathic helix so that it binds GTP in place of GDP (Behnia, Panic, Whyte, & Munro, 2004). This change allows ARF1 to bind the membrane where it plays an important role in recruiting coat complex protein I, clathrin adapter proteins AP1 and AP2, and other proteins necessary for proper vesicle budding and trafficking. This membrane binding is reversed by GTPase-activating proteins (GAPs) that catalyze hydrolysis of the GTP resulting in ARF1 release from the membrane. The nearly identical protein ARF3 has similar function, behaving indistinguishably in most studies. Given the importance of ARFs and the Golgi to cellular processes, bacteria often manipulate ARF function and Golgi stability to promote bacterial infection.

In this study, we show the pro-form of the *V. vulnificus* MARTX effector MCF is activated by autoprocessing upon binding with Class I ARF1 and ARF3. Although MCF can also be stimulated by ARF4 to autoprocess, these proteins do not associate in cells. Notably, MCF can be stimulated to auto-cleave by either the full-length forms of ARF1 and ARF3 or the more stable Δ17 forms of the proteins. This demonstrates that binding of ARF to MCF does not require the first 17 aa that are essential *in vivo* for membrane association.

The protein-protein interaction of MCF with ARF1 was confirmed in cells. ARF1 transitions between a GDP- and GTP-bound state dictating its position on and off of the Golgi (Behnia et al., 2004). As MCF is more rapidly and more fully induced to auto-cleave by ARF1-GTP than by ARF1-GDP, MCF is probably preferentially recruited to the Golgi, where it binds ARF1-GTP. Indeed, a low level of active ARF3-GTP at the Golgi under conditions for these experiments may explain our difficulty confirming MCF binding to ARF3 in cells (Tsai et al., 1993).

The cell data also support that once ARFs stimulate autoprocessing, they do not continue to stably associate with MCF, unless the catalytic cysteine is inactivated. This is in contrast to studies using purified proteins that show ARF1 can bind to both catalytically active and inactive MCF. We originally surmised this was evidence that ARF1 was modified in cells by catalytically active MCF and then released, although extensive analysis found no cleavage or post-translational modification of ARF1 or ARF3. We therefore conclude that there is a secondary event that alters the structure of MCF after processing to release ARFs and this exchange may require Cys-148.

Catalytically inactive cMCF was by contrast able to function as a protein trap for ARFs. Yet, this protein does not cause obvious cytotoxicity when overexpressed in cells (Agarwal, Agarwal, et al., 2015; Agarwal, Zhu, et al., 2015). These data contradict potential models whereby MCF functions simply to sequester ARF from its normal function. Thus, we surmise there is another protein that may be a target for cleavage by catalytically active cleaved MCF. Attempts to identify this host co-factor protein using catalytically inactive MCF as a substrate trap/bait for the initial immunoprecipitation were unsuccessful. It is possible that this protein may be cleaved or degraded in intoxicated cells and thus was difficult to identify using immunoprecipitation strategies.

Subsequent to autoprocessing, MCF was found to gain the ability to bind lipid PtdInsP5 and to undergo acetylation at the newly exposed N-terminal glycine. Recombinant MCF stimulated to process *in vitro* was not blocked to Edman degradation, even using crude eukaryotic cell lysate as a source of the host factor (Agarwal, Agarwal, et al., 2015). This observation suggests that MCF could be acetylated only in vivo by host N-acetyltransferases when MCF is present within cells Indeed, the ARF related protein ARFRP1 is N-terminal acetylated by NatC for recruitment to the Golgi. In our case, we surmise the acetylation may aid retention of MCF at the membrane following initial recruitment by ARFs. This retention may be augmented also by binding to PtdInsP5.

Our electron microscopy and immunofluorescence studies reveal that expression of active MCF in cells results in dramatic destruction of the Golgi and vesiculation of the cells. The destruction of the Golgi occurred also during natural delivery of the effector from bacteria. Unlike traditional IF techniques, using the novel cryoAPEX approach, the localization of MCF in cells could be successfully visualized. Although possible that acetylation and binding PtdInsP5 initially enriched MCF at the Golgi, the overexpressed protein is broadly distributed. It is found throughout the cell, enriched also on mitochondrial membranes and possibly the nuclear membrane. This is supported by cell fractionation that show MCF present in cytosol, nuclear, membrane, and mitochondria fractions (Supplemental Fig 6). At late time points, there were many darkly stained vesicles in the cell that might contain MCF although these are not easily distinguished by the cryoAPEX method from dark staining lysosomes. The direct association with mitochondrial membranes could explain the loss of the mitochondrial membrane potential that ultimately leads to mitochondrial mediated apoptosis through activation of Caspases. Likewise, Golgi dispersion is also known to induce apoptosis (Machamer, 2015). Thus, we identify the linkage of MCF to ARFs as the likely cellular process that results in MCF mediated cell death, while the precise molecular events subsequent to ARF stimulated activation of MCF remain to be resolved.

Other bacterial effectors have been identified as directly interacting with ARFs. For example, *Shigella flexneri* Type III secretion effector IpaJ is a member of the C58 family of cysteine peptidases but unlike MCF its activity depends on the canonical CHD active site. Furthermore, IpaJ does not autoprocess but instead, it directly recognizes and cleaves the myristoylated glycine of the lipid modified N-terminus of ARF1 and ARL GTPases. Similar to MCF, IpaJ preferentially targets the GTP-bound active form of ARF1. By cleaving ARF1, IpaJ releases the GTPase from the membrane, thus inducing Golgi disruption (Burnaevskiy et al., 2013). However, despite extensive effort, our negative data support that MCF does not likewise directly cause any processing or modification to ARFs as its mechanism for Golgi dispersion.

As noted above, ARFs are also allosteric activators of the ADP-ribosylating activity of cholera toxin and related heat-labile toxins from enterotoxigenic *Escherichia coli* (Moss & Vaughan, 1991). Similar to findings here, the N-terminal first 17 aa of ARF1 was dispensable for activation of cholera toxin (Hong, Haun, Tsai, Moss, & Vaughan, 1994). The structure of cholera toxin bound to ARF6 ultimately revealed that binding of ARF6 via its Switch 1 and Switch 2 regions to face of CTA1 ∼15 Å away from the CTA1 active site. Binding of ARF6 caused a structural remodeling of the CTA1 loop regions to allow nicotinamide adenine dinucleotide to bind to the active site for use to ADP-ribosylate the alpha subunit of the stimulatory G protein G_s_ (O’Neal, Jobling, Holmes, & Hol, 2005). Thus, there is precedence in *Vibrios* for binding of ARFs as an activator to stimulate structural remodeling of a bacterial effector, then without ARF for the effector to function on a different target protein. Similar structural studies of MCF and MCF bound to ARFs could elucidate these mechanisms for MCF.

These similarities support that stimulation by ARFs is a unifying theme for *Vibrio* toxins. Although most *V. vulnificus* MARTX toxins have MCF, the MARTX toxin for Biotype 3 strains lack this effector (Ziolo et al., 2014). However, these toxins do delivery a distinct MARTX effector we designated DomainX (DmX). DmX is similar in size to MCF and likewise shows homology to the C58 cysteine peptidases. However, DmX is only 22% identical to MCF and lacks the signature RCD/Y motif. Further, the aligned His and Asp catalytic residues shared with the C58 peptidases were found to be essential for DmX, whereas they were dispensable in MCF (Agarwal, Agarwal, et al., 2015; Kim & Satchell, 2016). DmX was likewise found to autoprocess upon stimulation by ARF1, ARF3, and ARF4; however, all three ARFs bound consistently with DmX in cells, unlike MCF which shows a preference for ARF1. Expression of DmX in cells likewise results in dispersion of the Golgi (Kim & Satchell, 2016). However, unlike MCF, DmX does not have an N-terminus blocked to Edman degradation and is not thought to be post-translationally modified in cells. In addition, catalytically inactive DmX is cytotoxic when expressed in cells, suggesting ARF sequestering may play a more significant role in its cytotoxic mechanism. We postulate that these two MARTX effectors have an evolutionarily shared mechanism for stimulated autoprocessing to cause Golgi dispersion, but these two effectors have diverged to target the Golgi by distinct molecular mechanism. Still, the presence of either MCF or DmX in nearly all *V. vulnificus* MARTX toxins reveals that dissolution of vesicular trafficking is important for *V. vulnificus* pathogenesis.

## 4. Materials and Methods

### 4.1 Tissue Culture

Cos7 and HEK293T (generously provided by Dr. Richard Longnecker at Northwestern) cells were prepared and cultured in Dulbecco’s modified Eagle’s medium (DMEM) supplemented with 10% heat-inactivated fetal bovine serum (FBS) and 1% penicillin-streptomycin at 37°C in 5% CO_2_. Cells were seeded in 150 mm tissue culture dishes or 12-well tissue culture plates, and grown to 60 - 70% confluence.

### 4.2 Bacterial strains and plasmids

gBlocks containing Δ17 ARF1, Δ17 ARF3, Δ17 ARF4, Δ17 ARF6, ARF1, ARF3, MCF, MCF^C148A^, cMCF, and cMCF^C148A^ were synthesized by Integrated DNA Technologies (IDT) (Supplemental Table 1). These fragments were cloned into SspI digested vector pMCSG7 using Gibson assembly (New England Biolabs E2611L) (Antic, Biancucci, Zhu, Gius, & Satchell, 2015). FLAG-MCF^C148A^-HA, FLAG-cMCF^C148A^-HA, FLAG-MCF-HA, FLAG-cMCF-HA, and MCF-eGFP plasmids were generated as described in Agarwal *et al*. 2015.

ARF-Myc isoforms were constructed as in Kim and Satchell 2016. CMCP6 *rtxA1::bla* and CMCP6 *rtxA1::mcf-bla V. vulnificus* strains were generated as detailed in Agarwal *et al*. 2015. MCF-Apex2 was made by Gibson assembly using an APEX2 gBlock synthesized by IDT (Supplemental Table 1) cloned into the FLAG-MCF-HA expression plasmid. GST-Δ17 ARF1 was generously provided by Dr. Heike Folsch as described in Shteyn *et al*. 2011.

### 4.3 Transfections and Western blotting

For ectopic gene expression, cells were transfected using 1:3 µg of plasmid DNA to µl of 1 mg/mL polyethylenimine. After 4 hours, the media was exchanged with fresh media. Following an additional 14 hours, the cells were washed with phosphate buffered saline (PBS) and collected. Whole cells were lysed in lysis buffer (300 mM NaCl, 20 mM Tris-HCl pH 8, 1.1% Triton X-100) supplemented with ethylenediaminetetraacetic acid (EDTA) free protease inhibitor tablets (Thermo Scientific A32965) incubated for an hour at 4°C and centrifuged to remove cell debris. 40 µg of total protein in the whole cell lysate was boiled for five minutes in 2× Laemmli sample buffer and separated by SDS-PAGE, followed by transfer to nitrocellulose membrane with the BioRad Mini TransBlot System. Western blots were then completed using anti-hemagglutinin (HA) (Sigma H6908), anti-Myc (Thermo Scientific PA1-981), anti-MCF (produced with purified MCF protein by Lampire Biological Laboratories, Pipersville PA), anti-ARF1 (Novus Biologicals NBP1-97935), anti-FLAG (Sigma F3165), anti-mouse Immunoglobulin G (IgG) (Li-Cor 926-32210 or 926-68070), or anti-rabbit IgG (Li-Cor 926-32211 or 926-68071) antibodies and visualized using the Li-Cor Odyssey Fc imaging system.

### 4.4 Immunoprecipitation

600 µg of total protein of Whole cell lysate was incubated overnight at 4°C with mouse IgG agarose (Sigma A0919), agarose removed by centrifugation, and lysate subsequently incubated overnight with EZview™ Red Anti-c-Myc Affinity Gel or EZview™ Red Anti-HA Affinity Gel (Sigma E6779 and E6654). Beads were washed with lysis buffer supplemented with EDTA-free protease inhibitor tablets four times, once with 10 mM Tris-HCl pH 8.0, and protein eluted with 2 M NaSCN in 50 mM Tris-HCl pH 8.0, 150 mM NaCl.

### 4.5 Protein production and *in vitro* cleavage assays

Plasmids were expressed in BL21 (DE3) *E. coli* (Wu, Christendat, Dharamsi, & Pai, 2000) to an OD_600_ of 0.8 in terrific broth, induced with 1 M isopropyl β-D-1-thiogalactopyranoside and grown overnight at 28°C. Cells were pelleted, lysed with a sonicator in Buffer A (10 mM Tris-HCl pH 8.0, 500 mM NaCl) with lysozyme, and cell debris removed by centrifugation. Δ17 ARF6 was recovered from the insoluble fraction by incubation for one hour in Buffer A with 6 M Urea, followed by centrifugation. Protein was then purified using the AKTA protein purification system with a Ni-NTA HisTrap affinity column (GE Healthcare GE17-5248-02), followed by passage through a Superdex 75 column for size exclusion chromatography. GST-Δ17 ARF1 was grown and lysate collected as above and batch purified using glutathione sepharose (GE 17-0756-01).

Purified proteins were used for cleavage assays in which 2.5 µM of specified MCF and 4 µM ARF isoform were incubated in RAX buffer (20 mM Tris pH 7.5, 150 mM NaCl, 10 mM MgCl_2_) for various time points at 37°C, Laemmli sample buffer added, and boiled for five minutes prior to separation by SDS-PAGE.

### 4.6 Bacterial intoxication

For bacterial treatments, mid-log phase CMCP6 *rtxA1::bla* and CMCP6 *rtxA1::mcf-bla V. vulnificus* strains were pelleted and resuspended in PBS. Cos7 cells were washed twice with PBS, treated with CMCP6 *rtxA1::bla* or CMCP6 *rtxA1::mcf-bla* at a MOI of 5 in DMEM, centrifuged at 500 × *g* for three minutes, and incubated for an hour at 37°C with 5% CO_2_.

### 4.7 Fluorescence Microscopy

For immunofluorescence microscopy, images were taken at the Northwestern University Center for Advanced Microscopy using the Nikon A1R+ GaAsP Confocal Laser Microscope or the Nikon N-SIM Structured Illumination Super-resolution Microscope.

Following treatment, cells grown on glass coverslips were washed with PBS and fixed with 4% formaldehyde for five minutes and permeabilized with 0.2% Tween 20. Cells were then blocked with 10% goat serum in staining buffer (1% bovine serum albumin (BSA) plus 0.05% Tween 20 in PBS) for one hour prior to staining with mouse anti-GM130 (BD Transduction Laboratories 610822) or mouse anti-Calreticulin (abcam ab22683) diluted 1:200 in staining buffer overnight. The coverslips were then washed five times for five minutes each time with staining buffer. Secondary goat anti-mouse Alexa Fluor 647 antibody (Thermo Scientific) was then added at 1:400 in staining buffer and incubated for one hour. The slides were washed twice with PBS and twice with double distilled water with two minutes between each wash. The slides were mounted using ProLong diamond antifade mountant with DAPI (4′,6′-diamidino-2-phenylindole) (Invitrogen P36962).

For live cell imaging, cells were grown on 35/10 mm glass bottom tissue culture dishes (Greiner Bio-One 627870). Prior to bacterial intoxication, cells were washed with PBS and incubated with 150 nM MitoTracker™ Green FM (Thermo Fisher M7514) in DMEM for 45 minutes. The cells were then washed with PBS and incubated with 5 µg/mL Hoescht 33342 (Immunochemistry Technologies 639) in PBS. After 10 minutes the cells were washed, fresh DMEM added, and treated with bacteria as previously described. Images were taken every 5 minutes following intoxication for an hour.

### 4.8 Electron microscopy and CryoAPEX localization

HeLa cells were grown in 10 cm dishes and transfected with MCF-Apex2 using Lipofectamine 3000 (Thermo Fisher). Cells were trypsinized 10 or 15 hours post-transfection, resuspended in DMEM and then pelleted at 500 *x* g. For live HPF cells were centrifuged and resuspended in 20% BSA in PBS. Cells were pelleted and loaded onto copper membrane carriers (1mm x 0.5 mm; Ted Pella Inc.) and cryofixed using the EM PACT2 high pressure freezer (Leica). Cryofixed cells were processed by FS using an AFS2 automated FS unit (Leica) using the extended FS protocol as described in Sengupta *et al*. 2019. For preparation of samples for cryoAPEX localization, cells were instead resuspended in 0.1% sodium cacodylate buffer containing 2% glutaraldehyde for 30 minutes following trypsinization, washed three times with 0.1% sodium cacodylate buffer and once with cacodylate buffer containing 1mg/ml 3,3’-DAB (Sigma-Aldrich). Pellets were then incubated 20 minutes in 1mg/ml DAB (Sigma) and 5.88mM hydrogen peroxide (EM sciences) in cacodylate buffer, pelleted and washed twice for 5 minutes in cacodylate buffer, once with DMEM, once with DMEM 20% BSA, and pelleted again. These cells were loaded onto copper membrane carriers and cryofixed as described before. An extended FS protocol was optimized for the preferential osmication of the peroxidase-DAB byproduct to ascertain membrane association of MCF. Briefly, frozen pellets were incubated for 60 hours at 90°C in acetone containing 5% water and 1% osmium tetroxide then at −20°C for a 6 hour osmication cycle. For FS of direct HPF samples, pellets were incubated for 24 hours at −90°C in acetone containing 0.2% tannic acid, washed three times for 5 minutes with glass distilled acetone (EM Sciences), and resuspended in acetone containing 5% water, 1% osmium tetroxide (OS) and 2% uranyl acetate (UA). Resin exchange was carried out by infiltrating the sample with a gradually increasing concentration of Durcupan ACM resin (Sigma-Aldrich) as follows: 2%, 4% and 8% for 2 hours each and then 15%, 30%, 60%, 90%, 100% and 100% + component C for 4 hours each. Samples were then embedded in resin blocks and polymerized at 60°C for 36 hours. Post-hardening, planchets were extracted by dabbing liquid nitrogen on the membrane carriers. Blocks were embedded a second time and hardened at 60°C for 36 hours. Serial sections, 90 nm thick were obtained using a Leica UC7 microtome. Serial-section ribbons were collected on formvar coated copper slot grids (EM sciences). Sample grids from HPF processing were stained for an hour with 4% UA solution in water followed by Sato’s Lead. CryoAPEX samples were not post stained. Screening was carried out using Tecnai-T12 transmission electron microscope operating at 80kV and images collected with a Gatan 4K camera.

### 4.9 Lipid overlay assays

PIP Strip™ Membranes (Thermo Fisher P23750) were probed as per the manufacturer instructions. Briefly, the membrane was blocked for an hour using TBS-T (10 mM Tris-HCl, pH 8.0, 150 mM NaCl, 0.1% Tween-20) + 3% fatty acid-free BSA (Sigma 126609) (TBS-T + BSA) for an hour at room temperature. The His-tag was removed from purified MCF and cMCF proteins using Tobacco Etch Virus (TEV) protease. Membranes were subsequently incubated with 0.5 µg/mL of either of these proteins in TBS-T + BSA overnight at 4°C and washed 10 times with TBS-T + BSA. The membrane was then reprobed and washed as before. Bound protein was detected using anti-MCF and anti-rabbit IgG antibodies as per the western blotting protocol.

### 4.10 Two Dimensional Western analysis

Anti-HA pulldowns of FLAG-MCF^C148A^-HA, FLAG-cMCF^C148A^-HA, FLAG-MCF-HA, and FLAG-cMCF-HA recovered as described above were prepped using the Bio-Rad ReadyPrep 2-D Cleanup Kit (Bio-Rad 1632130). The sample was then loaded onto 11 cm pH 3-10 immobilized pH gradient (IPG) strips (Bio-Rad 1632014) and rehydrated overnight. Isoelectric focusing was run on a Bio-Rad Protean IEF cell as per the Bio-Rad recommendations. Following IEF, the IPG strip was equilibrated and analyzed by SDS-PAGE and western blotting.

### 4.11 Mass Spectrometry Analysis

Protein embedded in gel bands was digested with 200 ng trypsin (Thermo Scientific) at 37°C overnight. Digest products were extracted from the gel bands in 50% acetonitrile/49.9% water/0.1% trifluoroacetic acid (TFA) and desalted with C18 StageTips prior to analysis by tandem mass spectrometry. Peptides were separated on an EASY-Spray HPLC column (25 cm × 75 µm ID, PepMap RSLC C18, 2 µm, Thermo Scientific) using a gradient of 5 - 35% acetonitrile in 0.1% formic acid and a flow rate of 300 nL/min for 45 minutes delivered by an EASY-nLC 1000 nUPLC (Thermo Scientific). Tandem mass spectra were acquired in a data-dependent manner with an Orbitrap Q Exactive mass spectrometer (Thermo Fisher Scientific) interfaced to a nanoelectrospray ionization source.

The raw MS/MS data were converted into MGF format by Thermo Proteome Discoverer (VER. 1.4, Thermo Scientific). The MGF files were then analyzed by a MASCOT sequence database search. The search was performed using fixed modifications of Met oxidation and Cys carbamidomethylation and variable modifications of Lys acetylation and N-terminal acetylation. Mass accuracy of 10 ppm on precursor ions and 15 ppm of fragment ions were allotted. Peptides identified by the MASCOT search were then manually validated.

To determine the activator of MCF, pulldowns completed on FLAG-cMCF^C148A^-HA, ectopically expressed in HEK293T cells were analyzed for total protein identification by mass spectrometry at the University of Illinois at Chicago Mass Spectrometry Core.

## ACKNOWLEDGEMENTS

The authors thank Heike Folsch for sharing ARF expression plasmids. We thank the Purdue CryoEM Facility at Hockmeyer Hall for access to the high-pressure freezer, freeze substitution unit, microtome and chemical hood. We thank Dr. Christopher Gilpin for access to the T12 electron microscope at the Purdue Life Sciences EM Core Facility where thin sectioning was carried out. We also thank Dr. Arvanitis and the Center for Advanced Microscopy and Nikon Imaging Center at Northwestern University for their invaluable help and resources. This work was supported by National Institutes of Health grants AI092825 (to K.J.F.S), R01GM10092 (to S.M.), as well as Indiana Clinical and Translational Sciences Institute grant CTSI-106564 (to S.M.), and Purdue University Institute for Inflammation, Immunology and Infectious Diseases grant PI4D-209263 (to S.M.), GM104610 (to R.R.O.L.), and GM103479 (to J.A.L.). A.H. was supported by a Ruth L. Kirschstein Institutional National Research Service Award (5T32AI007476). J.M. was supported by a UCLA Molecular Biology Institute Whitcome Fellowship. Immunoflourescence imaging work was performed at the Northwestern University Center for Advanced Microscopy generously supported by NCI CCSG P30 CA060553 awarded to the Robert H Lurie Comprehensive Cancer Center.

## SUPPORTING INFORMATION

Additional supporting information for this file includes six supplemental figures and one supplemental table.

**Supplemental Figure 1**. MCF does not cleave or directly modify ARF1 or ARF3.

A-G. Bottom-up mass spectrometry was performed on ARF1 and ARF3 samples recovered from recombinant (A) (B) or anti-Myc IPs (C-G) from HEK293T cells transfected or incubated with and without MCF or MCF^CA^. Modifications that were detected are denoted. Any modifications found on either ARF1 or ARF3 were not reproducible across replicates. Furthermore, There were no modifications detected that were found in ARF + MCF samples that were not found in ARF samples alone and thus not attributable to MCF.

**Supplemental Figure 2**. Edman degradation is blocked in MCF recovered from cells.

A. Reads from Edman degradation analysis completed on dt-MCF immunoprecipitated from HEK293T cell lysate using anti-HA antibody. No signal and very little background was detected.

B. Gel of sample used for analysis.

**Supplemental Figure 3**. MCF is N-terminally acetylated inside host cells.

Replicates of two-dimensional gel analysis of dt-MCF expressed in HEK293T cells, recovered by anti-HA immunoprecipitation. Amino acids acetylated in each circled population indicated in corresponding color on sequence.

**Supplemental Figure 4**. MCF does not alter normal endoplasmic reticulum structure.

Cos7 cells were transfected with empty vector or MCF-EGFP (green) for 18 hours, fixed, and stained for DAPI (blue), and endoplasmic reticulum marker (calreticulin) (red).

**Supplemental Figure 5**. Transmission electron microscopy shows Golgi vesiculation induced by MCF.

A, B. Electron microscopy tomograms taken of HeLa cells ectopically expressing MCF for 10 (A) or 15 hours (B). A-D. On right, higher magnification of boxed region on left showing Golgi (G), herniated Golgi (red arrows), vesicles (V). and autolysomes (A).

**Supplemental Figure 6**. Western blot of subcellular fractions from HEK 293T cells ectopically expressing dt-MCF. Each fraction probed with standards for membrane (CD-44), mitochondria (Hsp-60), and nucleus (HistoneH3), and MCF (anti-HA).

**Supplemental Table 1**. Sequences of gBlocks used for plasmid construction

